# Unbiased intestinal single cell transcriptomics reveals previously uncharacterized enteric nervous system populations in larval zebrafish

**DOI:** 10.1101/2022.08.11.503619

**Authors:** L.E. Kuil, N. Kakiailatu, J.D. Windster, E. Bindels, J.T.M. Zink, G. van der Zee, R.M.W. Hofstra, I.T. Shepherd, V. Melotte, M.M. Alves

## Abstract

The enteric nervous system (ENS) regulates many gastrointestinal functions including peristalsis, immune regulation and uptake of nutrients. Defects in the ENS can lead to severe enteric neuropathies such as Hirschsprung disease (HSCR), which is caused by defective ENS development. Zebrafish have proven to be fruitful in the identification of novel genes involved in ENS development and HSCR pathology. However, the composition and specification of enteric neurons and glial subtypes of the larval zebrafish at a single cell level, remains mainly unexplored. Here, we performed single cell RNA sequencing of zebrafish ENS at 5 days post-fertilization. We identified both vagal neural crest progenitors and Schwann cell precursors, as well as four clusters of early differentiated neurons. Interestingly, since we took an unbiased approach where we sequenced total intestines, an *elavl3+/phox2bb-* population of neurons and the presence of *cx43+*/*phox2bb-* enteric glia were identified in larval zebrafish. These populations have not been described before. Pseudotime analysis supported binary neurogenic branching of ENS differentiation, which happens via a notch-responsive state. Together, our data revealed previously unrecognized ENS populations and serve as a resource to gain new insights on ENS development and specification, proving that the zebrafish is a valuable model organism in the quest towards understanding and treating congenital enteric neuropathies.

## Introduction

The enteric nervous system (ENS) consists of neurons and glia that are tightly interconnected together, and with cells in their microenvironment. The function of the ENS extends far beyond regulating peristalsis, as it is also involved in secretion, immune regulation and nutrient absorption, via connections with other cell types in the intestine (1). It is well known that dysregulation of ENS development leads to life-threatening congenital enteric neuropathies, of which Hirschsprung disease (HSCR) is the most common disorder, affecting approximately 1 in 5.000 live births (1, 2). ENS development occurs early during embryogenesis with vagal and sacral neural crest contributions. Recently, it has been found in mice that at postnatal stages, the ENS is supplemented by enteric neurons derived from Schwann cell precursors (SCPs) (3). This suggests that there is a dual origin of ENS cells, namely those derived from embryonic (vagal) neural crest during early gut colonization, and those derived postnatally from SCPs (3).

One of the vertebrate animal models that is regularly used to study ENS development, is the zebrafish (4). Zebrafish are highly suitable for genetic manipulation, develop rapidly, ex-utero and are transparent, which makes them extremely valuable for screening novel disease genes and tracing developmental processes (5, 6). However, the precise composition and specification of different neuronal and glial subtypes in the zebrafish ENS, remains unclear. This holds true particularly at larval stages, in which key processes take place to ensure proper gut colonization with enteric neurons and glia. To date, a few immunohistochemistry studies investigating enteric neuronal identities in larval zebrafish, have reported the presence of vasoactive intestinal peptide (VIP), pituitary adenylate cyclase-activating peptide (PACAP), neuronal nitric oxide synthase (nNOS), serotonin (5-hydroxytryptamine; 5HT), calretinin (CR) and calbindin (CB), from 3 days post fertilization (dpf) onwards (7, 8). Recently, it has also been described that the adult zebrafish intestine contains enteric glia, similar to that observed in mamals (9). The study showed the presence of enteric glia presenting with neurogenic properties in the adult intestine, which could be detected by the notch reporter line *her4*.*3:*GFP (9). However, the existence of enteric glia in larval zebrafish, is still a controversial subject. Three papers reported contradicting findings regarding expression of canonical glial genes such as *gfap*, the traditional marker for enteric glia in human and mouse. Baker *et al*. showed Gfap+ enteric glia in the outer layer of the intestine of 7 and 18 dpf fish, encapsulated by a layer of enteric neurons (10). Transmission electron microscopy showed the presence of granular vesicles and filiform processes wrapping the muscularis and caveolae, which are typical characteristic of glia (10). McCallum *et al*. also showed Gfap+ staining in the larval intestine, but suggested that the immunostaining was aspecific, since it remained in the intestine of the *ret* mutant HSCR model, which lacks an ENS (9). Moreover, they showed that other typical enteric glial genes were not expressed in the zebrafish intestine, including *bfabp (fabp7a), sox10* and *s100b* (9). Such findings were supported by El-Nachef and Bronner, who reported the absence of enteric glia expressing *sox10, gfap, plp1a* and *s100b* in larval stages (11).

To gain new insights into the exact ENS composition of larval zebrafish, studies at the single cell transcriptome level are warranted. Previous zebrafish single cell transcriptomic studies described, did not capture enough neuronal cells for sub-analysis, or were done at very young embryonic and larval stages, showing limited neuronal specification (12-14). Here, we report single cell RNA sequencing (scRNA-seq) of 5 dpf zebrafish intestines. Importantly, we used an unbiased approach, dissecting whole intestines and sequencing all live cells without enrichment for specific ENS markers, such as *sox10* and/or *phox2bb*. Such approach allowed detection of previously unrecognized neuronal and glial populations in the larval intestine, expanding our understanding of the ENS composition and specification in larval zebrafish.

## Results

### Vagal derived ENS cells are complemented by Schwann cell precursors (SCPs), supporting the dual origin of the ENS in zebrafish

To enable capturing of the ENS from 5-day-old tg(*phox2bb*:GFP) larvae (15), 244 intestines were pooled to perform 10x scRNA-seq. Based on expression of canonical markers, as well as markers obtained in previous literature (16-18), we selected clusters that most likely contained neural crest progenitors, enteric neurons and glia (e.g. expression of *phox2bb, elval3/4, sox10, slc1a2b*) (n = 1369 cells; 15% of total cells) (Fig S1). Subset analysis of these cells, led to eleven distinct clusters (Fig 1A, 1B). Two of these clusters were characterized by shared expression of typical neural crest markers such as, *sox10, foxd3* and *phox2bb* and were therefore, classified as progenitor cells (Fig S2A). However, while one cluster selectively expressed genes typical for oligodendrocyte precursor cells (OPCs) or Schwann cell precursors (SCPs), including *clic6, tppp3*, and *anxa1a* (16, 19-22) (n = 50 cells; Fig 1B, S2B), the other cluster showed specific expression of more traditional (vagal) neural crest genes, such as *ret, hoxb5b, a*nd *tlx2* (23-26) (n = 181 cells; Fig 1B, S2C). *Mmp17b*, which has been described in migrating trunk neural crest and in Schwann cells upon injury, was specifically identified in the SCP cluster (Fig S2B) (20, 27, 28). We then performed single molecule fluorescent whole mount in-situ hybridization (smFISH) using a probe targeting *mmp17b*, to localize these cells in 5 dpf zebrafish and determine if they are specifically present in the gut. As expected, positive cells were present in the spinal cord and in the axonal motor neuron branches, corresponding to the known localization of SCPs (Fig 1C) (20, 29, 30). In the intestine, *mmp17b* signal was also observed, occasionally co-localizing with the *tg(phox2bb:*GFP) signal (Fig 1D). This signal was sparse, which is in line with our scRNA-seq data, where the majority of *mmp17b* positive cells (32 out of 52 cells) showed only 1 or 2 RNA counts/cell. Therfore, our results confirm the presence of SCPs in the zebrafish intestine, and support the rare nature of these cells at 5dpf.

**Figure 1.**
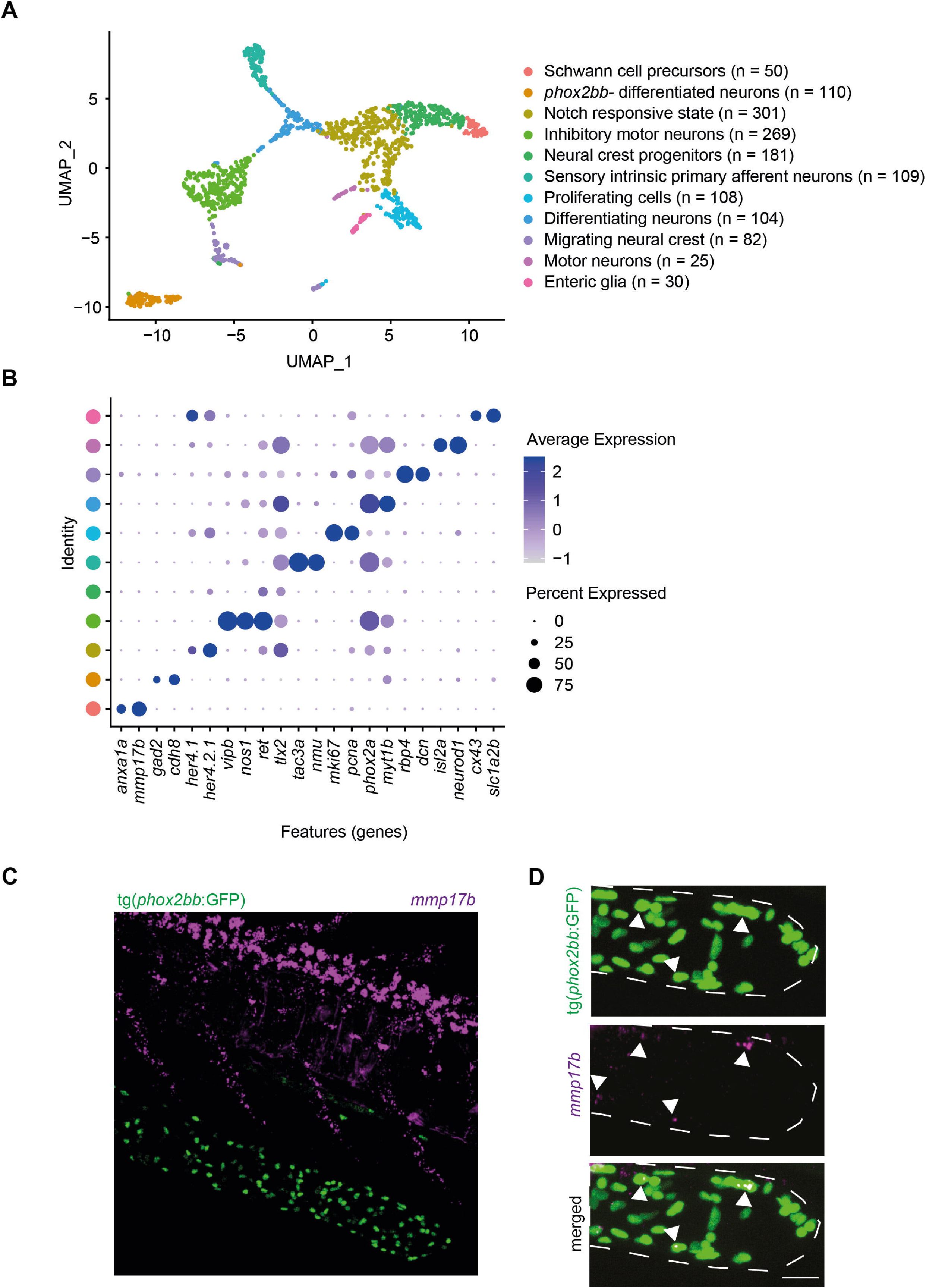
Single cell transcriptomics of 5 dpf zebrafish ENS. A) UMAP of 1369 ENS cells, containing eleven different clusters. B) Dot plot showing expression of genes highly differentially expressed between clusters. C) smFISH of 5 dpf tg(phox2bb:GFP) larvae stained for mmp17b (magenta). Scale bar represents 50 μm. D) Zoom images of smFISH of 5 dpf tg(phox2bb:GFP) larvae showing co-localization of mmp17b (magenta) and phox2bb (green) in the intestine, outlined with dotted lines. Arrows highlight cells of interest showing colocalization. Scale bar represents 10 μm.

### The zebrafish intestine contains four types of differentiated neurons at larval stage

Based on our analysis, four types of ‘differentiated neurons’ were identified. The largest cluster (n = 269 cells) consisted of inhibitory motor neurons, expressing *vip* and *nos1* (Fig 1B, 2A) (31, 32). This cluster also included cells expressing *slc6a4b, tph1b* and *ddc*, which are genes involved in serotonin transport and production (Fig S3D-F) (33-35). Sensory intrinsic primary afferent neurons (IPANs) were identified by expression of *nmu, vgf, tac3a* and *calb2a* (n = 109 cells; Fig 1B, 2A, S3A) (32, 36). A third cluster expressing *isl2a/b, olig2* and *neurod1*, most likely represents motor neurons (n = 25 cells; Fig 1B, 2A, S3B) (37-39). The fourth cluster seems to contain a mix of different neuronal subtypes such as, glutamatergic, GABAergic and IPANs (n = 110 cells), based on their selective expression of *vglut2a (slc17a6b), gad1b, sv2a, neurod6, gad2* and *cdh8* (Fig 2A, 2B, S3C) (32, 40, 41). Interestingly, this latter cluster did not express the neural crest marker *phox2bb* (Fig 2B).

**Figure 2.**
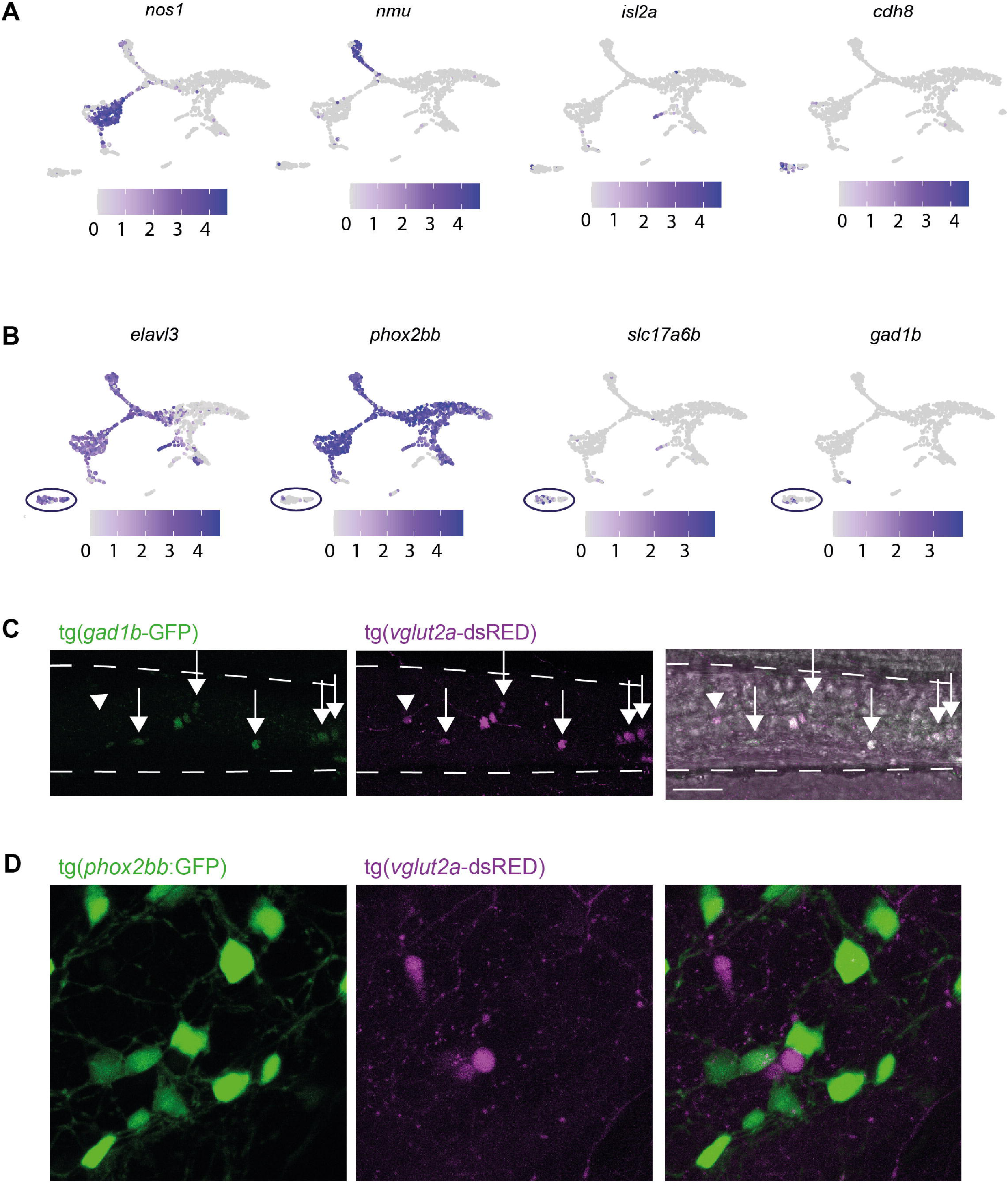
Cluster of elavl3+;phox2bb-enteric neurons present with excitatory and inhibitiory gene expression. A) Featureplots of genes defining four clusters of differentiated neurons. B) Featureplots highlighting the presence of a cluster of cells expressing elavl3, slc17a6b and gad1b, but lacking expression of phox2bb (phox2bb-differentiated neurons, depicted by the circle). C) Live imaging of 7 dpf tg(gad1b:GFP);tg(vglut2a-dsRed) larval intestine shows overlap between the two reporters (arrows) with one gad1b+ cell that is vlgut2-(arrowhead). Scale bar represents 40 μm. D) Live imaging of 7 dpf tg(phox2bb:GFP);tg(vglut2a-dsRed) larval intestine shows no overlap between the two reporters. Scale bar represents 10 μm.

### Identification of a *phox2bb-* population of differentiated enteric neurons

To confirm the presence of *phox2bb*-neurons in the zebrafish intestine, we first showed co-localization of the tg(*vglut2*:loxp-dsRed-loxP-GFP) and the tg(*gad1b*:GFP) reporter lines in the intestine (Fig 2C). Subsequently, we crossed the tg(*phox2bb*:GFP) reporter with the tg(*vglut2*:loxp-dsRed-loxP-GFP) reporter and found no co-localization between *phox2bb* and *vglut2 in vivo*, confirming the presence of *phox2bb-*/*vglut2a+* cells in the zebrafish intestine (Fig 2D). Our transcriptomic data also showed that these cells express *elavl3* (encoding HuC; Fig 2B), which led us to perform a HuC/D staining on 5 dpf tg(*phox2bb*:GFP) fish. We observed that a limited number (between 0 to 25) of HuC/D+;*phox2bb*-cells were present in the zebrafish intestine, comprising on average 2.5% of the total ENS (Fig 3A). This number, roughly corresponds to our scRNA-seq data (8% of total ENS cells). Distribution of HuC/D+;*phox2bb*-cells seems equal along the anterior to posterior axis, indicating that these cells are evenly distributed along the total length of the intestine and therefore, do not seem to be region specific (Fig 3A).

**Figure 3.**
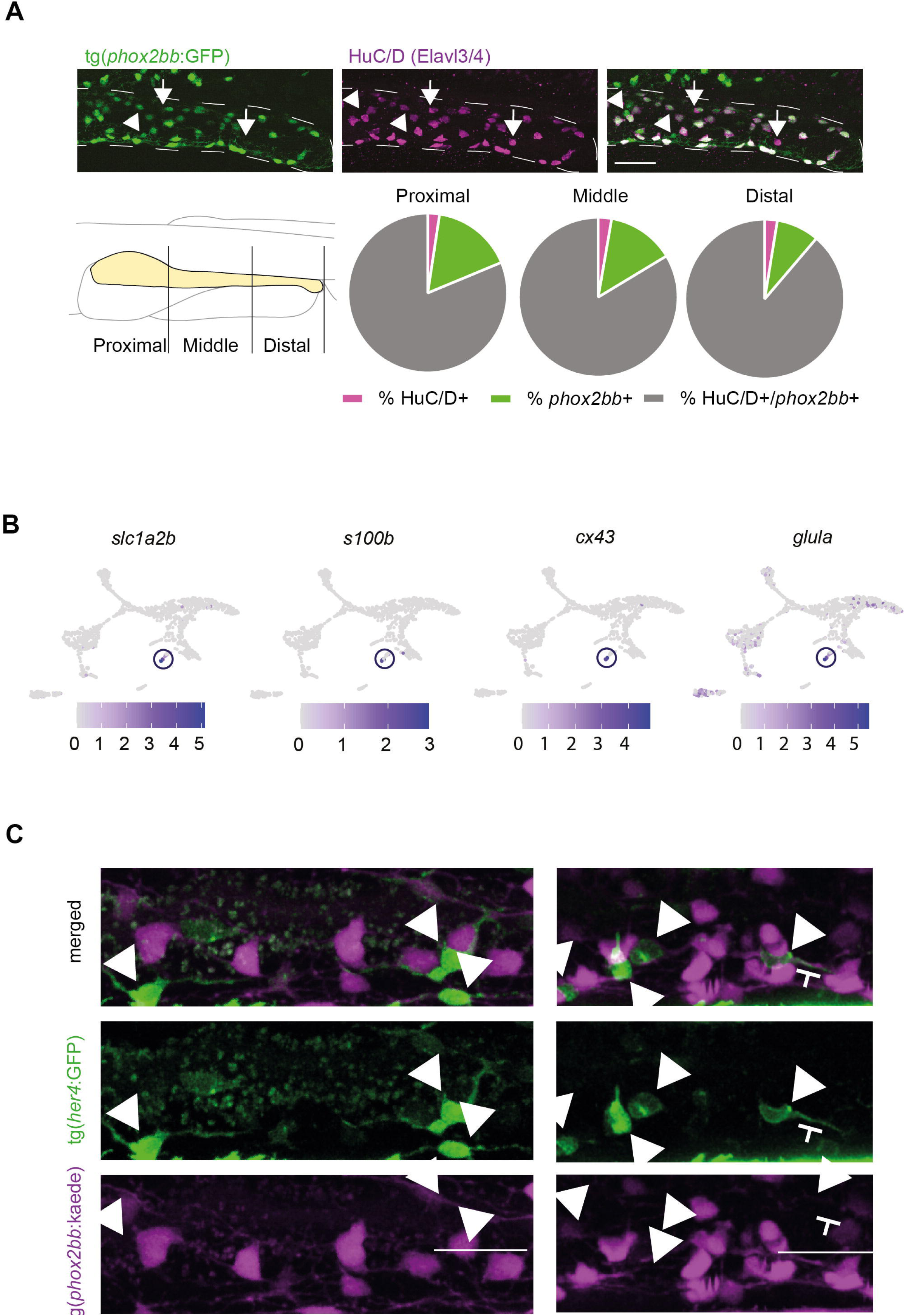
One small cluster expresses genes typical for enteric glia in mammalians. A) HuC/D antibody staining shows that most HuC/D+ cells in the intestine express phox2bb, but also show phox2bb+/HuC/D-cells (progenitors) depicted by arrowheads and phox2bb-;HuC/D+ cells (differentiated neurons) depicted by arrows. Scale bar represents 40 μm. Quantification of the relative amount of double and single positive cells are presented in pie charts (n=9). B) Featureplots showing selective expression of cx43, glula, slc1a2b and s100b in one specific cluster of enteric glia depicted by the circle. C) Live-imaging of 5 dpf tg(8.3phox2bb:kaede);tg(her4:GFP) intestines shows phox2bb-;her4+ cell depicted by the arrowheads that are in close proximity to, or seem to interact with phox2bb+ neurons. Scale bar represents 20 μm.

### Larval zebrafish intestine contains enteric glia

Although recent compelling evidence suggest the presence of enteric glia in the adult zebrafish intestine (9), previous studies are unambiguous about the existence of these cells in larval zebrafish. This is mainly due to the absence of expression markers typical for this cell type in mice and humans. In line with previous reports, our data confirmed the absence of *gfap+* cells in the larval intestine of the tg(*gfap*:GFP) reporter line (Fig S4C). However, due to our unbiased approach of sequencing total intestines, we were able to observe a cluster of cells lacking expression of *phox2bb* and *sox10*, but expressing Hairy/E(spl)-related 4 (*her4*) genes (n = 30 cells; Fig S4A). Interestingly, this cluster showed highly specific expression of genes typically found in radial glia in the zebrafish brain, such as *glula, slc1a2b* and *ptn* (16), and of genes expressed in mammalian enteric glia, such as *cx43, s100b, sox2, ptprz1b*, and *fabp7a* (Fig 3B, S4B) (42-46). Analysis of tg(*her4:*GFP);tg(*phox2bb*:kaede) larvae showed that *her4+*;*phox2bb*-cells are indeed present in the intestine and are located in close proximity to, or in some cases in direct contact to, *phox2bb*+ cells (Fig 3C). To validate the enteric glial identity of these cells, we performed immunohistochemistry on tg(*phox2bb:*GFP) larvae at 3, 4, 6 and 10 dpf using an antibody against connexin 43 (Cx43), a known enteric glia marker in mice that we found expressed in the Cx43+/*phox2bb-* cluster. Based on our results, Cx43+ cells were detected from 4 dpf onwards, suggesting that enteric glia arise between 3 and 4 dpf in zebrafish (Fig 4A). At 4 dpf, Cx43+ cells were most often observed in the middle intestinal segment, with an average of 12 cells per fish (Fig 4B). Although the location of Cx43+ cells is similar to *phox2bb*+ cells, as they were often observed in the same focal plane in close proximity to each other, these cells were always negative for the *phox2bb* reporter.

**Figure 4.**
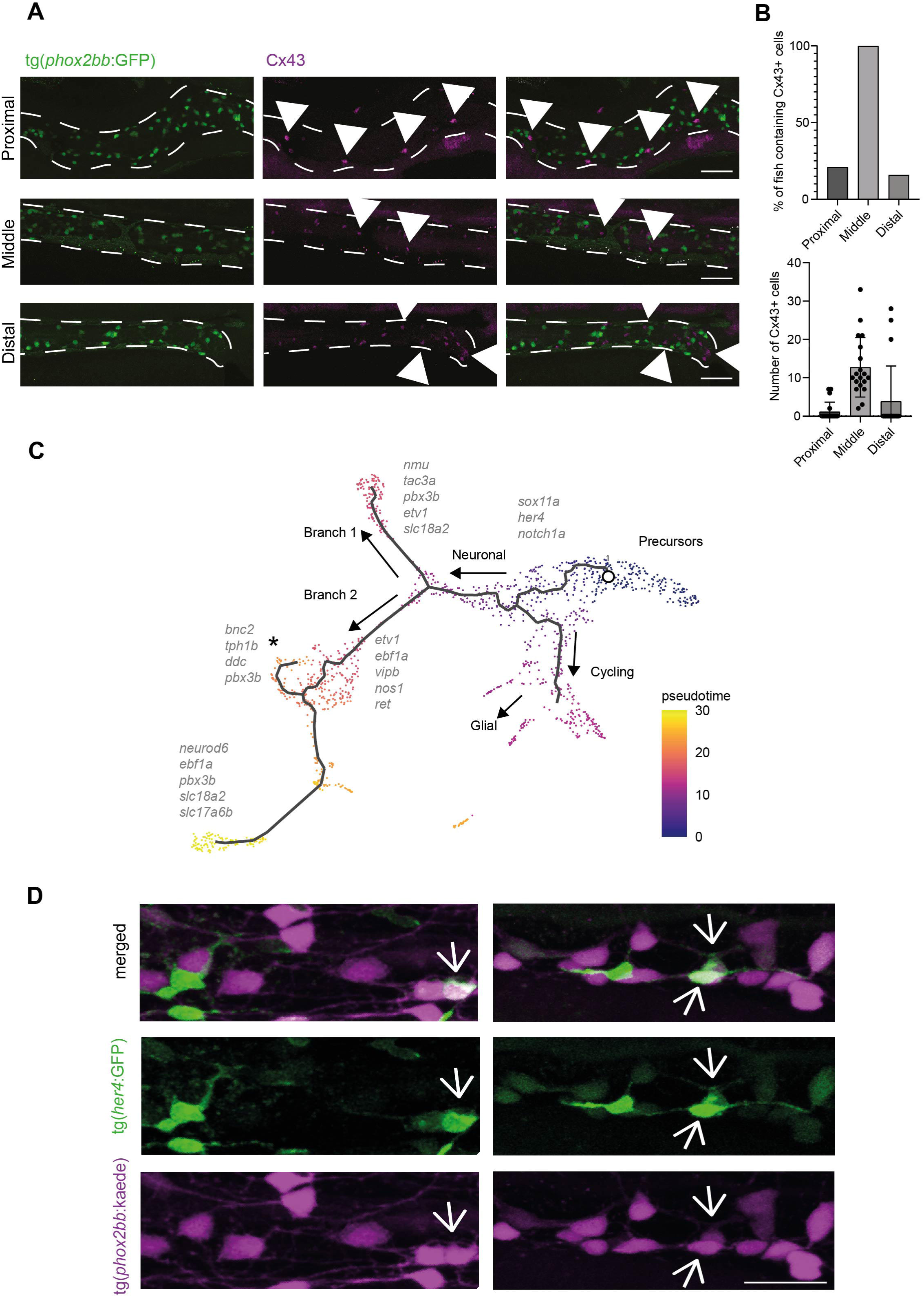
Pseudotime analysis shows differentiation trajectories from right to left of the UMAP. A) Immunohistochemistry staining of Cx43 in the tg(phox2bb:GFP) reporter line shows non overlapping expression in the intestine. Representative images from 4 dpf larvae. Scale bar represents 40 μm. B) Upper graph showing the percentage of larvae that contained Cx43 cells in their proximal, middle and distal intestine. The lower graph shows the number of Cx43 cells per larvae in the proximal, middle and distal intestine (n=19). C) Pseudotime color-coded featureplot showing a bifurcation towards neuronal differentiation (sensory IPAN: branch1 and inhibitory motor neurons: branch2 containing a secondary branch towards serotonergic neurons marked with the asterix). D) Live-imaging of 5 dpf tg(8.3phox2bb:kaede);tg(her4:GFP) intestines shows phox2bb+;her4+ cells, representing cells undergoing differentiation from progenitor state towards neuronal or glial fate. Scale bar represents 20 μm.

### Progenitor cells become notch-responsive before differentiation towards neuronal and glial fate

RNA expression of a marker for differentiated neurons, *elavl3*, and a gene specific for early neural crest/progenitor cells, *sox10*, showed that these genes are expressed exclusively on the left and right side of the UMAP, respectively. Analysis of key cell fate mediators for pan-neurogenic fate (e.g. *elavl4* and *insm1b)* or genes expressed in newborn neurons in the zebrafish brain (e.g. *tubb5* and *tmsb*), showed a similar pattern of expression on the left side of the UMAP (Fig S5A) (16). This suggests a trajectory of progenitors’ differentiation towards neurons, from right to left in the UMAP. Pseudotime analysis by monocle3 (47) confirmed this differentiation trajectory and additionally showed that during differentiation, a bifurcation into two types of early differentiated neurons occurs, branch 1: sensory IPANs versus branch 2: inhibitory motor neurons (Fig 4C). This latter branch also contains a secondary branch towards serotonergic neurons (see asterix in Fig. 4C). In line with this finding, Morarach et al. reported a similar bifurcation in the murine ENS differentiation trajectory, with branch A, forming Vip/Nos positive neurons (32). Expression of “branch A marker genes” *etv1* and *ebf1a*, was also found in our dataset, in the *vip+/nos1+* inhibitor motor neuron branch (Fig 4C, S5C). Conservation in expression of various genes at specific differentiation states is depicted in figure 4 panel C (32). Comparing our data to the dataset from Howard *et al*. showed that the early differentiation observed in zebrafish from 68-70 hours post fertilization (hpf), is prominently seen in our dataset (14). For example, co-expression of *slc18a2* and *pbx3b* was observed in differentiated neurons including IPANs (Fig 4C).

Our data also showed that neuronal differentiation seems to start in notch-responsive cells, specifically expressing *notch1a, notch3* and notch-responsive *her4* genes (Fig S5B) (9). In line, our UMAP showed co-expression of the known cell fate mediator gene *sox11a*, and notch (responsive) genes (Fig S5B) (16). Live-imaging of 5 dpf tg(*her4*:GFP)/tg(8.3*phox2bb*:keade) fish showed that a subset of *phox2bb*+ cells is indeed *her4*+ (15% of cells in the posterior intestine), confirming the presence of *phox2bb*+*/her4*+ notch responsive cells *in vivo* (Fig 4D). Thus, progenitor cells seem to become notch responsive upon initiation of differentiation towards neurons (branch 1 and 2). In addition, there is a third trajectory emanating progenitors towards clusters containing proliferative cells, motor neurons and enteric glia, suggesting a separate differentiation route towards a cycling/enteric glial fate (Fig 4C).

## Discussion

Here, we present for the first time a single cell atlas of the ENS of 5 dpf zebrafish. Our results show the presence of two clusters of progenitor cells, traditional vagal neural crest cells and Schwann cell precursors (SCPs), confirming a dual origin of the zebrafish ENS. Based on our dataset, we were able to identify *mmp17b* as a specific marker for SCPs, and were able to confirm the presence of these cells in the gut, as well as their rare nature *in vivo*. Our data, also showed the presence of four clusters of enteric neurons and a cluster of enteric glia, at this developmental stage. Inhibitory motor neurons were confirmed to be major contributors of the zebrafish ENS (7, 8), but we were also able to identify other differentiated neuronal subtypes. We can clearly see that early enteric neuronal differentiation occurs as an initial bifurcation towards two major branches. Differentiation of sensory IPANs seem to develop via branch 1, whereas *vip+/nos1+* inhibitory motor neurons specify via branch 2. The latter, seems to contain a secondary branch towards serotonergic enteric neurons. Therefore, differentiation of enteric inhibitory motor neurons as well as serotonergic neurons, seems to be conserved between at least, mice and zebrafish (32). Interestingly, due to our unbiased sequencing approach in which the whole intestine was analysed, we were able to identify one cluster of *elavl3+*/*phox2bb*-differentiated neurons, expressing genes specific for glutamatergic neurons, GABAergic neurons, and others involved in serotonergic signaling. Based on our live imaging data, *elavl3+*/*phox2bb*-neurons are located in close proximity or sometimes even directly adjected, to *phox2bb+* enteric neurons. To our knowledge, such population has never been defined before, as all enteric neurons were assumed to express *phox2bb*. Future lineage tracing experiments should be performed to confirm the neural crest-origin of these cells, as well as scRNA-seq experiments at older ages, to provide insights into which neuronal sub-types these *phox2bb*-cells contribute.

Finally, our dataset showed the presence of a cluster of enteric glial cells. In line with previous studies, we found that the relative contribution of enteric glial to the ENS seems to be less abundant in zebrafish, compared to that in human and mice. We now show that enteric glia can be detected already at zebrafish larval stages. We also confirmed that canonical enteric glial genes such as, *sox10*, and *plp1a* are not expressed in the putative enteric glial cluster. However, we did detect RNA expression of *s100b* and *fabp7a*, which is in contrast to previous studies (9, 11). In addition, we showed that canonical glia in larval zebrafish express *cx43, notch3* and *her* genes, but lack *phox2bb* expression. The *her4*-reporter line previously described, has specifically characterized enteric glia in the adult zebrafish intestine (9). However, here we show that only the *phox2bb-*/*her4+* cells, but not the *phox2bb+/her4+* cells, express enteric glial markers at larval stages. Interestingly, *her4* expression was not only limited to enteric glia, but was also consistently found in *phox2bb*+ ‘intermediate cells’ undergoing differentiation. A previous study had already shown that enteric neural crest cells start to express *her4* after migration, and lose its expression upon differentiation (9). Here, we extended their findings by showing differentiation at a single cell transcriptional level, from progenitors via a notch responsive state, towards early specification of enteric neuronal fate. Together, this suggests that Notch signaling plays a central role in the transition from progenitor to neuronal state or glial differentiation. In line with this, disruption of Notch signaling in mice (*Pofut1* knockout), was shown to result in the absence of an ENS, confirming that this signaling pathway is crucial to maintain the neural crest progenitor pool (48). The Notch pathway has also been recognized in the maintenance of neuronal stem cells in the brain, but signaling dynamics in neuronal differentiation have yet to be elucidated (Reviewed by (49)).

Taken together, our results show that the zebrafish ENS has a dual origin of precursor cells, that follow specification upon notch activation towards either a neuronal fate, or via a cycling state towards an enteric glial fate. It also shows that using an unbiased approach in which cells are not selected for a specific reporter construct, can be instrumental to find new cell clusters. In summary, our data adds to the understanding of healthy ENS development and offers an essential framework for intra-study, cross-species, and disease state comparisons.

## Acknowlegdements

We want to thank Remco Hoogenboezem for mapping the scRNA-seq data, Emma de Pater for useful discussions, and the optical imaging center (OIC) of the Erasmus University Medical Center for assistance with confocal microscopy.

## Funding

This work was funded by the friends of Sophia foundation (SSWO WAR-63).

## Supplementary figures

***Supplementary figure 1. Featureplots showing the clusters selected for subset analysis of the ENS***

*A) UMAP featureplots showing the cells that are selected for subset analysis of the ENS in purple and other non-ENS cells in grey. B) Featureplots showing expression of phox2bb, GFP, her4*.*1 and elavl3 in the ENS subset*.

***Supplementary figure 2. Featureplots showing specific gene expression in various clusters***

*A) UMAP featureplots showing expression of genes in the clusters containing A) precursors;*

*B) SCPs, as well as C) vagal neural crest cells and their derivatives*.

***Supplementary figure 3. Featureplots showing specific expression in differentiated neuronal clusters***

*UMAP featureplots showing expression of genes in the clusters containing A) sensory IPANs; B) motor neurons; C) phox2bb-neurons (sv2a+), and D) serotonergic neurons (in the inhibitory motor neurons cluster)*.

***Supplementary figure 4. Featureplots showing specific expression in enteric neurons***

*A) UMAP featureplots showing the absence of expression of phox2bb and sox10, but presence of her4 expression in the enteric glial cluster depicted by the circle. B) UMAP featureplots showing expression of genes in the enteric glial cluster, depicted by the circle*. *C)Live-imaging capture of the tg(gfap:GFP) reporter line showing the absence of GFP+ cells in the intestine, which is outlined by dotted lines. Abbreviation: AF = autofluorescence*.

***Supplementary figure 5. Featureplots showing expression of genes at specific differentiation states***

*A) UMAP featureplots showing expression of genes typical for cells committed to neuronal differentiation. B) UMAP featureplots showing expression of notch genes, notch responsive her4*.*2*.*1. and the cell fate mediator gene sox11a. C) UMAP featureplots showing expression of genes that define differentiation branches previously described (32)*.

## Material and methods

### Animal husbandry

The following zebrafish lines were used: transgenic tg(*phox2bb*:GFP)(15), tg(*8*.*3phox2bb:keade)*(50), tg(*her4*:GFP)(51), tg(gfap:GFP)(52), tg(*vglut2*:loxp-dsRed-loxP-GFP)(53), and tg(*gad1b*:GFP) (54). Zebrafish were kept on a 14/10h light/dark cycle. Embryos and larvae were kept in an incubator at 28.5°C in HEPES-buffered E3 medium. For imaging experiments, fish were treated from 24 hpf onwards, with 0.2 mM 1-phenyl 2-thiourea (PTU), to inhibit pigmentation. Animal experiments were approved by the Animal Experimentation Committee of the Erasmus MC, Rotterdam.

### Isolation of zebrafish intestines

Intestines of 5 days post-fertilization (dpf) larvae were isolated as followed: a row of 6-10 larvae anesthetized with 0,0016% Tricaine, were placed on an 1.8% agarose plate. Intestines were isolated using insect pins under a dissection microscope (Olympus SZX16), collected with a tweezer and placed in an Eppendorf tube containing phosphate buffered saline (PBS) with 10% fetal calf serum (FCS), on ice. In total 244 intestines were isolated and pooled together.

### Pre-processing of zebrafish cell suspension for scRNA sequencing

Cells were dissociated using 2.17mg/mL papain dissolved in HBSS, with CaCl_2_ and MgCl_2_. Papain was activated using 2.5µl cysteine (1M) and dissociation was performed in a water bath at 37°C, for 10 minutes. Cells were then transferred into a FACS tube using a 35 μm cell strainer. Cells were centrifuged at 700g for 5 minutes at 4°C, the supernatant was removed and pellets were resuspended in PBS containing 10% FCS. DAPI was added to mark dead cells (1:1000). All sorts were performed using the FACSAria III sorter, into eppendorfs containing PBS with 5% FCS.

### Single cell RNA sequencing (scRNA-seq)

Single cells were barcoded using a 10x genomics Chromium Controller, and sequenced using a Novaseq 6000 instrument (Illumina). In total, 9.858 cells were sequenced with mean reads per cell of 21.106. For scRNA-seq analysis we used Seurat V3 (55). The Seurat pipeline was used for filtering (nFeature_RNA > 100 & nFeature_RNA <4200 & percent.mito < 0.05), normalization and downstream analysis for clustering, where we used 50 dimensions with a resolution of 0.8 for the UMAP processing. This led to 49 clusters, which we annotated based on differential gene expression. Seven clusters expressing neuronal and/or enteric progenitor markers were identified (cluster 6, 10, 13, 19, 21, 23, 34)(Fig. S1).

These seven clusters were selected for a subset analysis, using 30 dimensions with a resolution of 0.7 for the clustering and UMAP, resulting in 14 clusters. One cluster was excluded as it contained leukocytes (*lcp1*+, *phox2bb-, elavl3-, sox10-*; cluster 12) and proliferating cells negative for *phox2bb* and *elavl3* were manually excluded. The final set was analyzed using six dimensions and a resolution of 0.4. This analysis provided us with eleven clusters, which were annotated based on the differential gene expression and literature search. If required, a more thorough analysis of the total set of differentially expressed genes per cluster, or differentially expressed genes between specific subclusters, was performed. For pseudotime analysis, monocle3 was used (47).

### Fluorescent imaging

Imaging was performed as previously described (56). For the keade photoconversion experiments, all *phox2bb+* cells in the total intestines were photoconverted using the 405 nanometer (nm) laser, as described earlier (11). After photoconversion, the green and red channels were recorded, using a sequential scan with the 488nm and 561nm lasers to confirm full photoconversion.

### Immunohistochemistry

Whole mount immunohistochemistry using mouse anti-HuC/D (1:100, molecular probes A-21271) was performed, as previously described (57). Antibody staining using rabbit anti-Cx43 (1:200, Cell Signaling Technologies 83649) was performed as published before (58). To increase signal to noise ratio we made use of monovalent AffiniPure Fab Fragments (111-167-003 Jackson; Cy™3 AffiniPure Fab Fragment Goat Anti-Rabbit IgG).

### Single-molecule whole-mount fluorescent in situ hybridization

Zebrafish were fixed in 4% paraformaldehyde in PBS, overnight. They were then dehydrated through a series of 25/50/75/100% MeOH in PBST for 5 minutes each, and stored for a minimum of 1 hour at −20°C. Next, samples were rehydrated through a series of 75/50/25/0% MeOH in PBST for 5 minutes each, and incubated in prot K for 15 minutes at 20 °C. They were rinsed twice with PBST for 5 minutes and re-fixed in 4% PFA in PBS, for 20 minutes at room temperature. Susequently, samples were rinsed again 5×5 minutes in PBST. After manual pre-treatment for permeabilization, we continued with the RNAscope Multiplex Fluorescence Reagent Kit v2 Assay (Advanced Cell Diagnostics, Bio-Techne), according to the manufacturers’ instructions. A custom made probe for dr-*mmp17b* C1 (NPR-0035110) was used (Advanced Cell Diagnostics, Bio-Techne). Opal 570 dye (Akoya Biosciences) was used for channel development.

## Notes

### Competing Interest Statement

The authors have declared no competing interest.

### Summary of Updates

Corrected the typo in the title

