## Supplementary figures and images for "Unbiased intestinal single cell transcriptomics reveals previously uncharacterized enteric nervous system populations in larval zebrafish"

### Fig S1

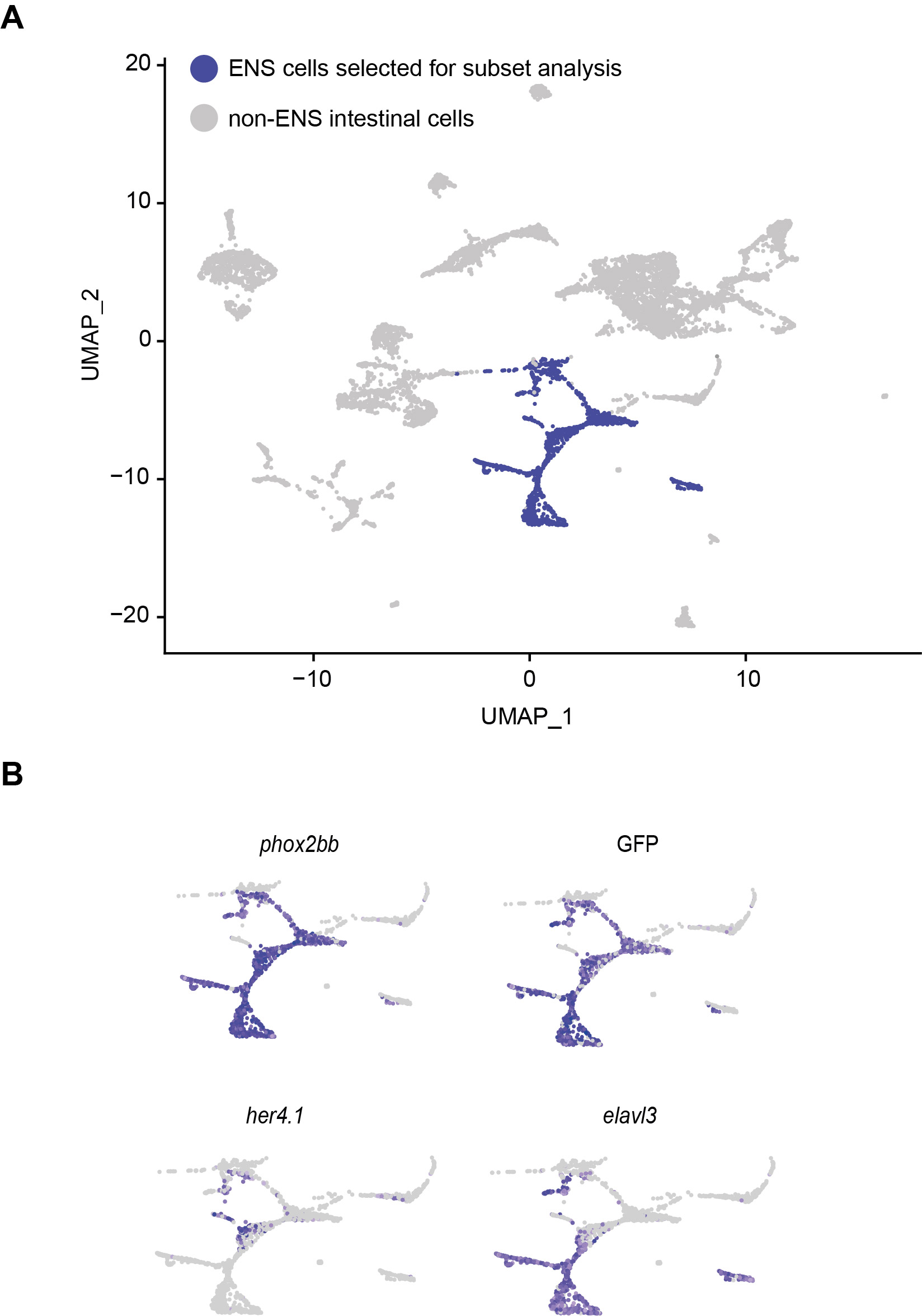

### Fig S2

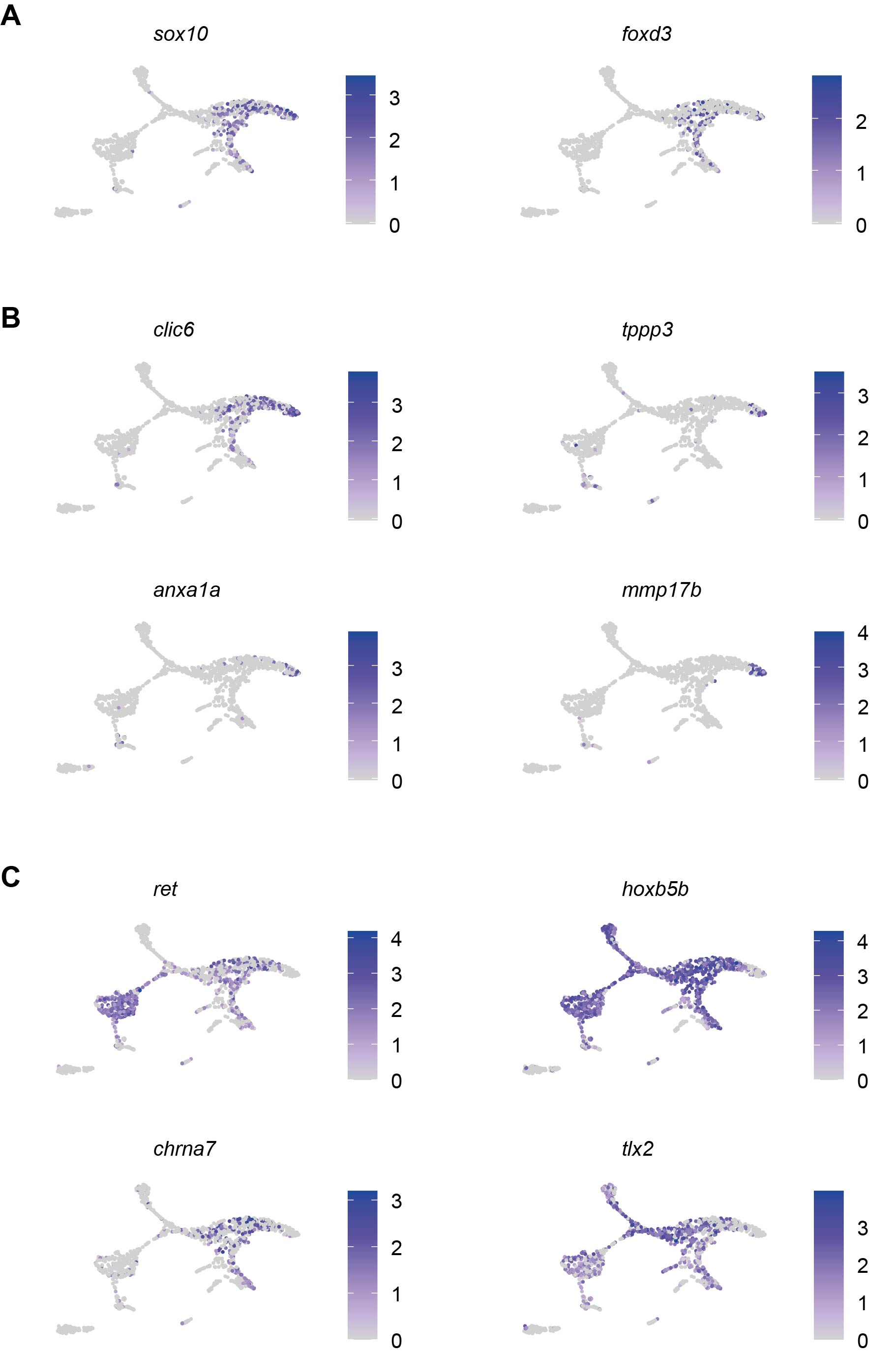

### Fig S3

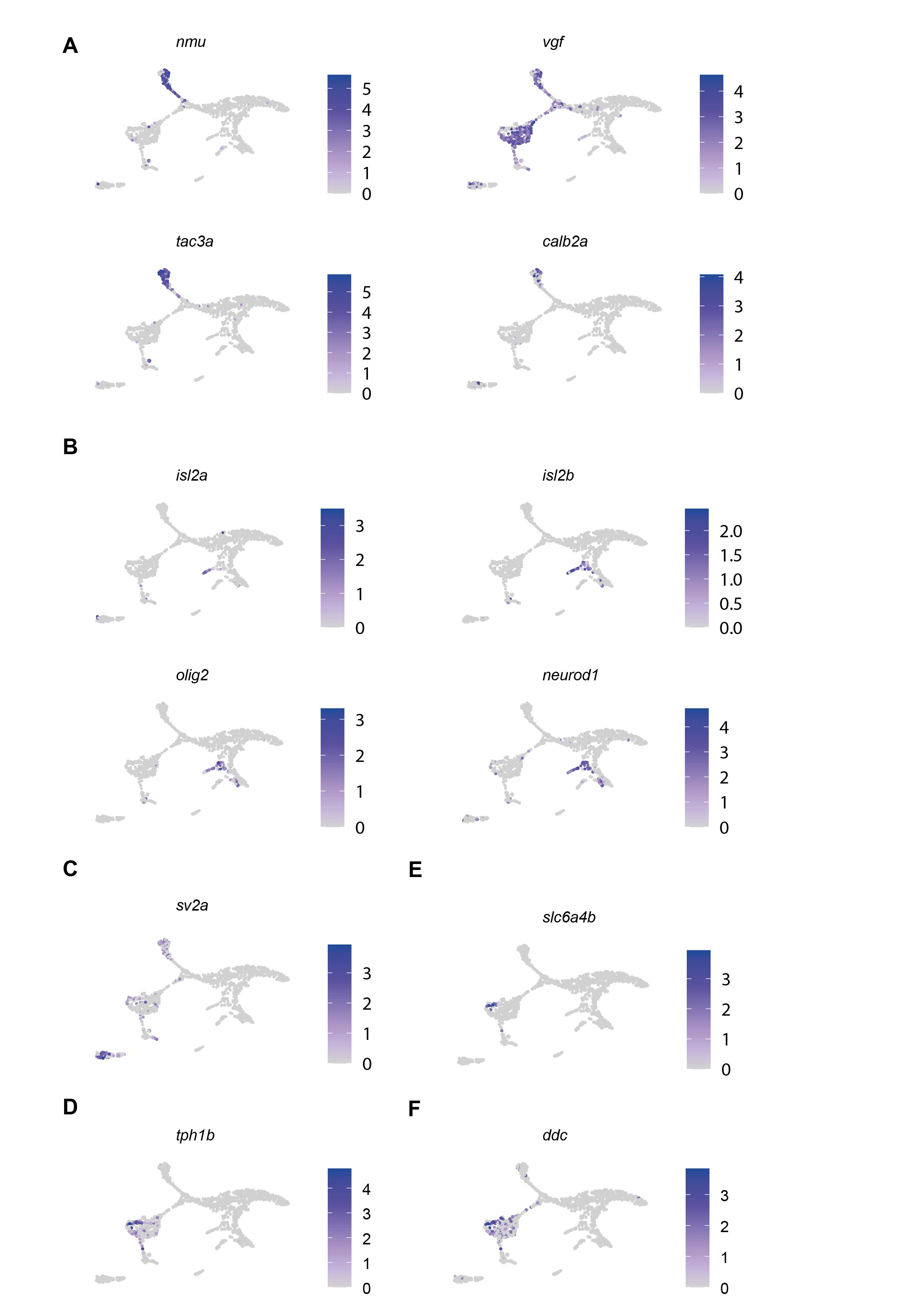

### Fig S4

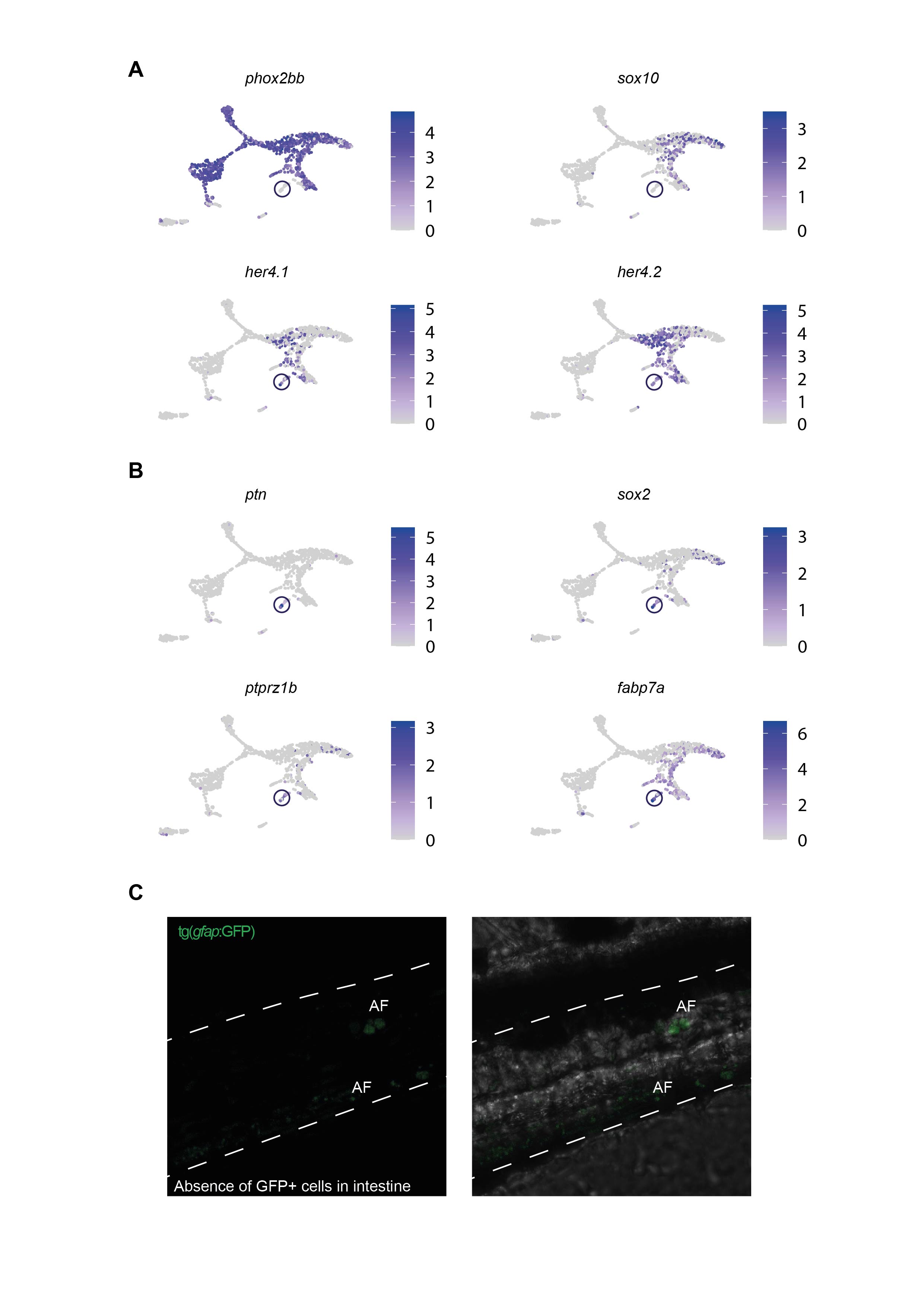

### Fig S5

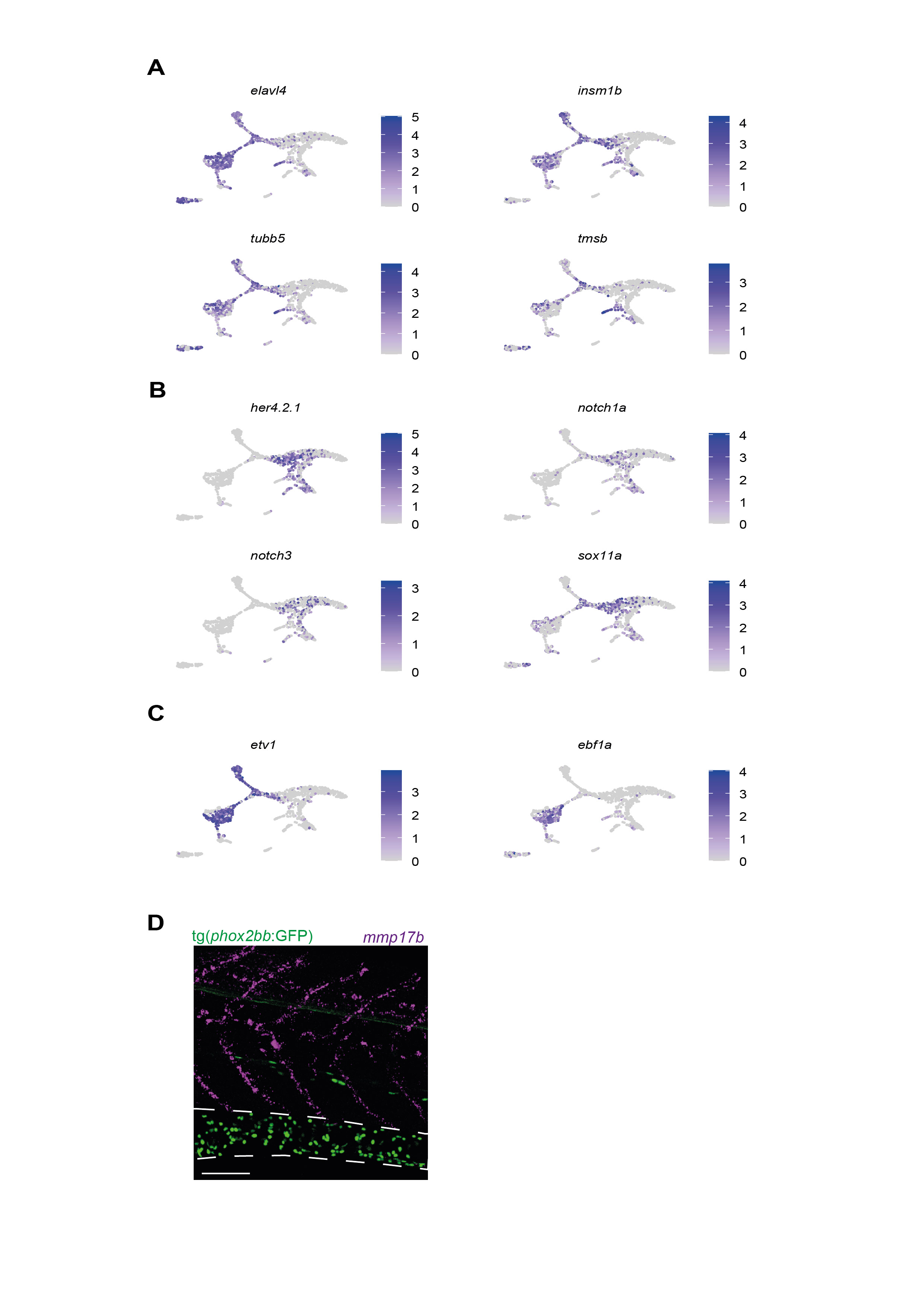
